# Acoustically Targeted Noninvasive Gene Therapy in Large Brain Regions

**DOI:** 10.1101/2023.01.19.524626

**Authors:** Shirin Nouraein, Sangsin Lee, Vidal A. Saenz, Huckie C. Del Mundo, Joycelyn Yiu, Jerzy O. Szablowski

**Author notes:** Correspondence should be addressed to J.O.S.

## Abstract

Focused Ultrasound Blood-Brain Barrier Opening (FUS-BBBO) can deliver adeno-associated viral vectors (AAVs) to treat genetic disorders of the brain. However, such disorders often affect large brain regions. Moreover, the applicability of FUS-BBBO in the treatment of brain-wide genetic disorders has not yet been evaluated. Herein, we evaluated the transduction efficiency and safety of opening up to 105 sites simultaneously. Increasing the number of targeted sites increased gene delivery efficiency at each site. We achieved transduction of up to 60% of brain cells with comparable efficiency in the majority of the brain regions. Furthermore, gene delivery with FUS-BBBO was safe even when all 105 sites were targeted simultaneously without negative effects on animal weight, neuronal loss, or astrocyte activation. To evaluate the application of multi-site FUS-BBBO for gene therapy, we used it for gene editing using the clustered regularly interspaced short palindromic repeats (CRISPR)/CRISPR-associated 9 (Cas9) system, and found effective gene editing, but also a loss of neurons at the targeted sites. Overall, this study provides a brain-wide map of transduction efficiency and the first example of gene editing after site-specific noninvasive gene delivery to a large brain region.

## INTRODUCTION

Brain disorders are often caused by dysfunction in multiple regions of the brain^1–3^. Gene therapy is a promising approach for treating many such disorders, including multiple sclerosis^4^ and genetic disorders such as Huntington’s^5^ or Tay–Sachs disease^6^. Recently, multiple clinically approved gene therapeutics and clinical trials^7^ have used adeno-associated viral vectors (AAVs), owing to their low pathogenicity, immunogenicity, and durable long-term gene expression^8^. However, delivery of AAVs to large brain regions is a major challenge. The presence of the blood-brain barrier (BBB) limits the uptake of most AAV serotypes from systemic circulation into the brain. Therefore, clinical trials rely on alternative delivery methods, such as local intraparenchymal injection^9^, which is efficient but requires multiple injections to cover the volume of even a single brain region^10^. Consequently, the majority of clinical trials focus on brain disorders that have focal pathogenesis, such as focal epilepsies ^11^, Parkinson’s^12^ disease, or Huntington’s disease^5^. Therefore gene delivery in the context of brain disorders involving multiple brain regions has to be enhanced.

Focused ultrasound (FUS) in combination with microbubble-induced BBB opening is a noninvasive^13,14^ method for AAV delivery to the brain. FUS-BBBO transiently opens the BBB to allow clinically used AAVs, such as AAV9, to enter the targeted regions in the brain from the blood^15^ with millimeter precision and without tissue damage^16–19^. Moreover, in principle, FUS-BBBO can also open the BBB throughout the brain^20^, providing widespread gene delivery, even for AAVs that cannot cross an intact BBB. If this is possible, FUS-BBBO could be used to treat disorders involving brain-wide pathophysiology, including inherited genetic disorders^21^. However, to date the FUS-BBBO transduction efficiency throughout the brain has not been rigorously evaluated.

Such disorders could also benefit from genome editing strategies such as clustered regularly interspaced short palindromic repeats (CRISPR) systems that use RNA-guided genome editing of DNA sequence^22–25^. CRISPR can edit mutations in vivo^26,27^, thus potentially treating inherited gain-of-function mutations^28^. To achieve this, CRISPR uses short RNA sequences, called single-guided RNAs (sgRNAs), which can direct a DNA nuclease, typically CRISPR-associated 9 (Cas9)^25,28^ to delete or modify target genomic sequences. For CRISPR to be useful in brain disorders, a feasible method of gene delivery to the brain is essential, which is currently not available. To date, approaches for CRISPR editing in the brain have used IC Injections of viral vectors^29^ and nanoparticles^30^; however, these approaches are not scalable to target large brain regions, unlike FUS-BBBO^31,32^.

To investigate the safety and feasibility of gene delivery to multiple brain regions using FUS-BBBO and to evaluate the transduction efficiency and cell-type tropism throughout the brain, we delivered AAV9 carrying various genetic cargo. First, we used AAV9 carrying green fluorescent protein (GFP) under a constitutive CaG promoter to target 11, 22, or 105 different brain sites with FUS-BBBO. Additionally, we tested whether a CRISPR/ Cas9 system could be effectively delivered in one AAV with FUS-BBBO and evaluated the transduction efficiency throughout the targeted brain regions. In summary, these experiments demonstrated that FUS-BBBO is safe and effective for non-invasive gene delivery throughout the brain without causing weight loss, neuronal loss, or other unfavorable outcomes and that gene editing with CRISPR/Cas9 shows variable efficiency when Cas9 is expressed under a promoter that fits within the AAV packaging capacity.

## RESULTS

### Safety and efficacy of FUS-BBBO targeted to multiple brain regions

Firstly, we examined the number of FUS-BBBO sites that could be targeted without affecting animal survival. Initially, we targeted 11 and 22 sites in one hemisphere and then attempted to target the largest volume of the brain that was technically feasible at a single depth of targeting with 85 sites. A separation distance was chosen to avoid overlap of the fullwidth half maximum (FWHM) pressure of the beam. The centers of each targeted site were separated by 1.5 millimeters with a FUS beam pointing downwards from the top of the brain (**Fig. 1**). We chose the parameters of opening based on previous studies^34,35^. Specifically, we chose 0.27 MPa of pressure, a frequency of 1.53 MHz, and a 10 ms pulse length repeated once per second for 30 s per site^36,37^. To rapidly test the feasibility and safety of the BBB opening parameters, we used FUS-BBBO to deliver the intravenous Evans blue dye (EBD). EBD is an easily detectable dye that is frequently used to evaluate the efficiency of FUS-BBBO using histology^38,39^. We targeted FUS-BBBO with EBD in nine male mice with a total of 363 opening sites. After 20 minutes we injected EBD intravenously and perfused the mice. We found that 100% of the sites showed successful delivery of the EBD, suggesting that the chosen parameters were useful for reliable BBB opening (**Supplementary Fig. 1**, n = 6 male mice, with 11, 25, or 85 sites opened per mouse, n = 2 per group). We also observed that EBD extravasation was specific to the targeted sites, suggesting that the EBD does not cross the BBB^39^.

**Figure 1.**
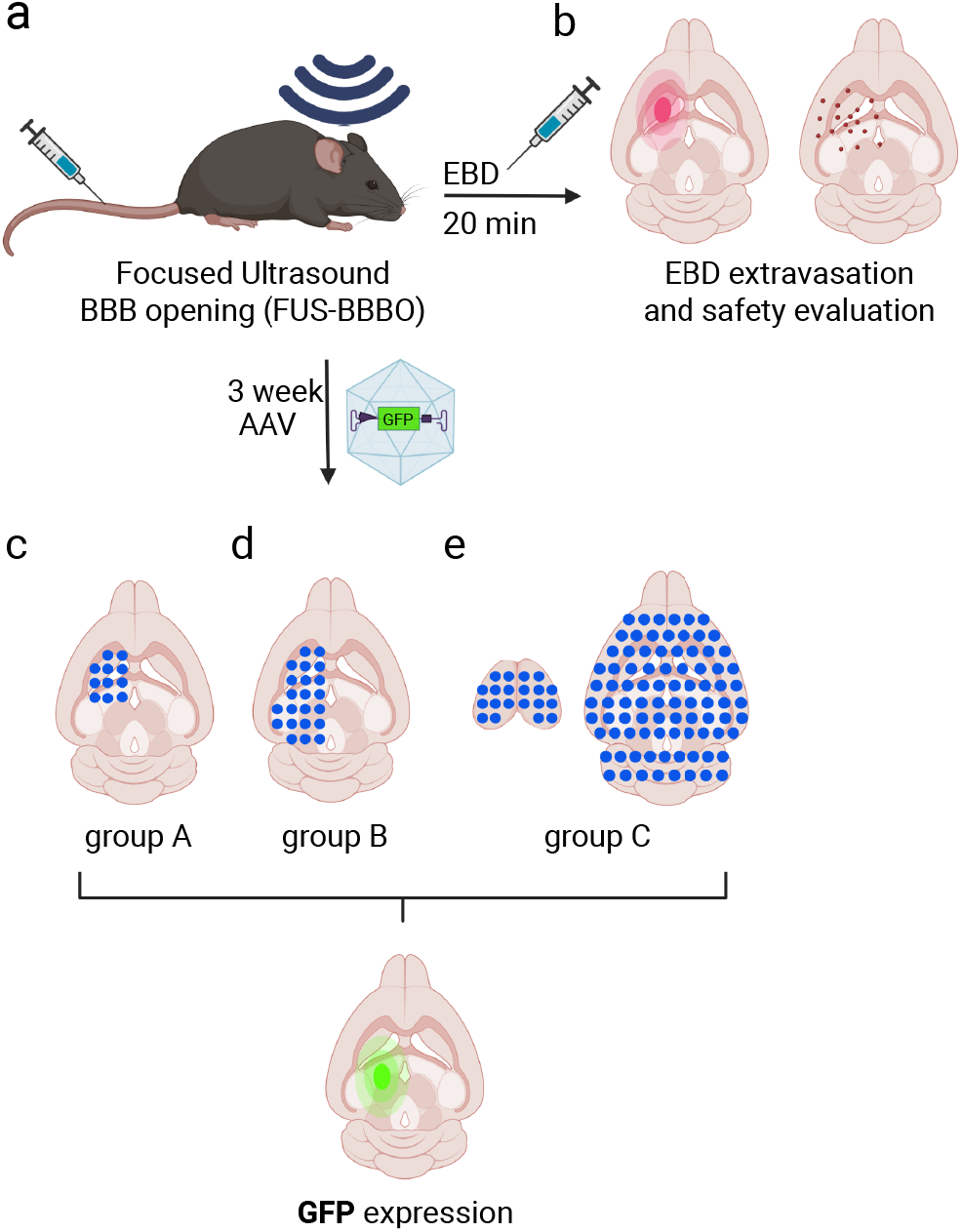
Achieving Effective & Safe FUS-BBBO over Large Brain Areas for Extensive AAV-mediated Gene Delivery. (a) Experimental overview for FUS optimization for (b) effective (successful EBD extravasation) and safe (minimal hemorrhagic occurrence) FUS-BBBO (C) successful AAVs carrying-GFP delivery under the CAG promoter with 0.27Mpa pressure, for 11 sites (group A, n=4) and (d) 22 sites (group B, n=5) and (e)105 sites (group C, n=6) showing the expression of the green fluorescent protein in all targeted site for three groups. (FUS-BBBO, Focused Ultrasound Blood-Brain Barrier Opening; adeno-associated viral vectors, AAV; Evans blue dye, EBD; green fluorescent protein, GFP). (Image created with Biorender.com)

Using the validated FUS-BBBO strategy and parameters, we tested the efficiency with which FUS-BBBO delivered AAVs into the targeted brain regions. We placed a nuclearly localized (NLS) GFP under the constitutive and broadly expressed CAG promoter (CAG-NLS-GFP). We packaged this plasmid in AAV9, which has been previously reported to be successfully delivered to the brain with FUS-BBBO^40^. We then injected 1E10 viral particles per gram of body weight, immediately followed by FUS-BBBO. Mice were sacrificed 3 weeks later, and brain sections (50 μm thickness) were stained against GFP to measure the transduction efficiency at each targeted site. We observed successful gene delivery at 100% of the targeted sites that could be recovered by tissue histology.

The percentage of transduction was analyzed within the FUS beam and was either compared to contralateral untargeted regions (groups A and B) or to a mouse that received an IV Injection of AAV, but no FUS-BBBO in group C (**Fig. 2a, b**). The sites were numbered for each group (**Supplementary Fig. 2**). The number of GFP-positive cells was significantly increased in targeted structures compared to that in untargeted sites. For 11-site targeting, the average transduction efficiency was 32.92±2.76%; for 22 sites, it was 44.99±2.55%, and for 105 sites targeting, it was 60.82±1.9% (p<0.0001 for each group, two-tailed t-test, paired for groups A and B, and unpaired for group C, **Fig. 2 c-e**). We observed that increasing the number of target sites significantly increased the transduction efficiency at each site, suggesting that widespread gene delivery with FUS-BBBO is an advantageous method for gene delivery (**Fig. 2f**). Transduction in untargeted sites did not vary between groups A and B, or A and C. However, there was a significant difference in the transduction of untargeted sites between groups B and C (p<0.05, one-way ANOVA with Tukey’s HSD test), possibly due to contralateral FUS-BBBO affecting transduction, as group B used contralateral controls, and group C used matched animals without FUS-BBBO (**Fig. 2g**).

**Fig. 2.**
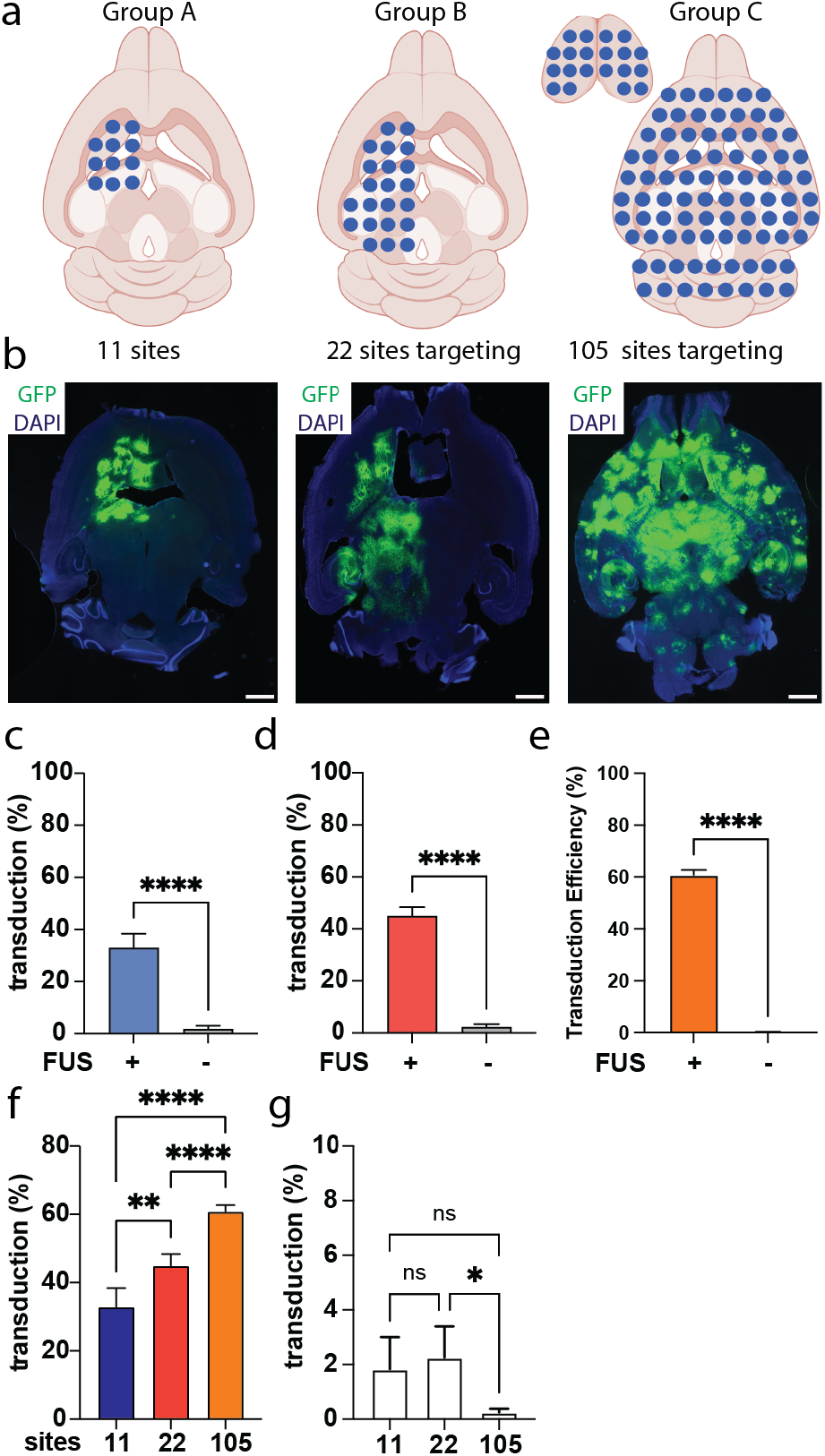
Gene delivery efficiency increases with the volume of BBB opened. (a) Blue dots indicate the FUS-BBBO delivery sites on axial drawings of the mouse brain. Immunohistochemistry of anti-GFP showed an increase in the number of GFP-positive cells in the brain of FUS-treated mice (b) representative brain sections from each group of mice shows the expression of the GFP in all targeted areas. c) For group A, GFP-positive cells were counted in the targeted area and the contralateral site showed higher efficacy of gene delivery after FUS-BBBO. The average transduction efficiency for each group was 32.92% for 11 sites at the targeted sites, and 1.8% at the contralateral sites, (p<0.0001, two-tailed heteroscedastic t-test; n=44 sites in 4 mice for each group). d) The average transduction efficiency in each targeted site for group B was 44.99% and 2.2% in contralateral sites (p<0.0001, two-tailed heteroscedastic t-test; 110 sites in 5 mice per group). e) Finally, for group C, the transduction efficiency was 60.82% at the targeted sites and 0.2% for the control mice that received AAV9 carrying GFP but no FUS-BBBO (478 sites in 5 mice for the targeted group, and 44 randomly selected sites in 3 control mice). f) The difference in transduction efficiency was significant between groups A-C, with a larger number of targets leading to a higher transduction per site [One-way ANOVA, F(2,629)=57.62, p<0.0001]. g) The transduction in untargeted sites ranged between 1.8% for 11 site-targeted groups, 2.2% for the 22-site-targeted group, and 0.24% for the 105-site group. There was a significant difference between the 22-site and 105-site groups (p<0.05, one-way ANOVA with Tukey’s post-hoc test). Notably, 11 and 22-site groups used contralateral controls, while 105 sites used brains from mice injected with AAV but without FUS-BBBO since the entirety of the brain was targeted preventing the use of internal contralateral controls. Scale bars are 1000 microns. *, p<0.05, **, p<0.01, ***, p<0.001, ****, p<0.0001. (FUS-BBBO, Focused Ultrasound Blood-Brain Barrier Opening; green fluorescent protein, GFP; Analysis of variance, ANOVA)

### Region-specific transduction efficiency after FUS-BBBO gene delivery

We then analyzed whether transduction efficiency varied between the brain regions and each of the 105 targeted sites after FUS-BBBO gene delivery. Each representative region is shown as a number in **Fig. 3a** and **Fig. 3b**. Transduction was successful at all analyzed sites. We observed transduction of 66.4±8.4% in the striatum (1), 69.5±12.6 % in the hippocampus (2), 54.5±4.4% in the cerebellum (3), 61.9±4.8% in the midbrain (4), and 60.1±8.9% in the cortex (5) (numbers are means with 95% CI, n=6 mice per group). The transduction efficiency was comparable between the two targeted hemispheres (58.4% vs. 57.3% on average, n=6 mice, p=0.76, two-tailed heteroscedastic unpaired t-test), indicating equally efficient delivery to each hemisphere. Representative images for each region are shown in **Fig. 3c-g**, with the top image showing the targeted site and the bottom image showing an analogous site from the control mice. There were no statistically significant differences between the groups (p=0.18, F(4,25)=1.713, One-way ANOVA, **Fig. 3f**).

**Figure 3.**
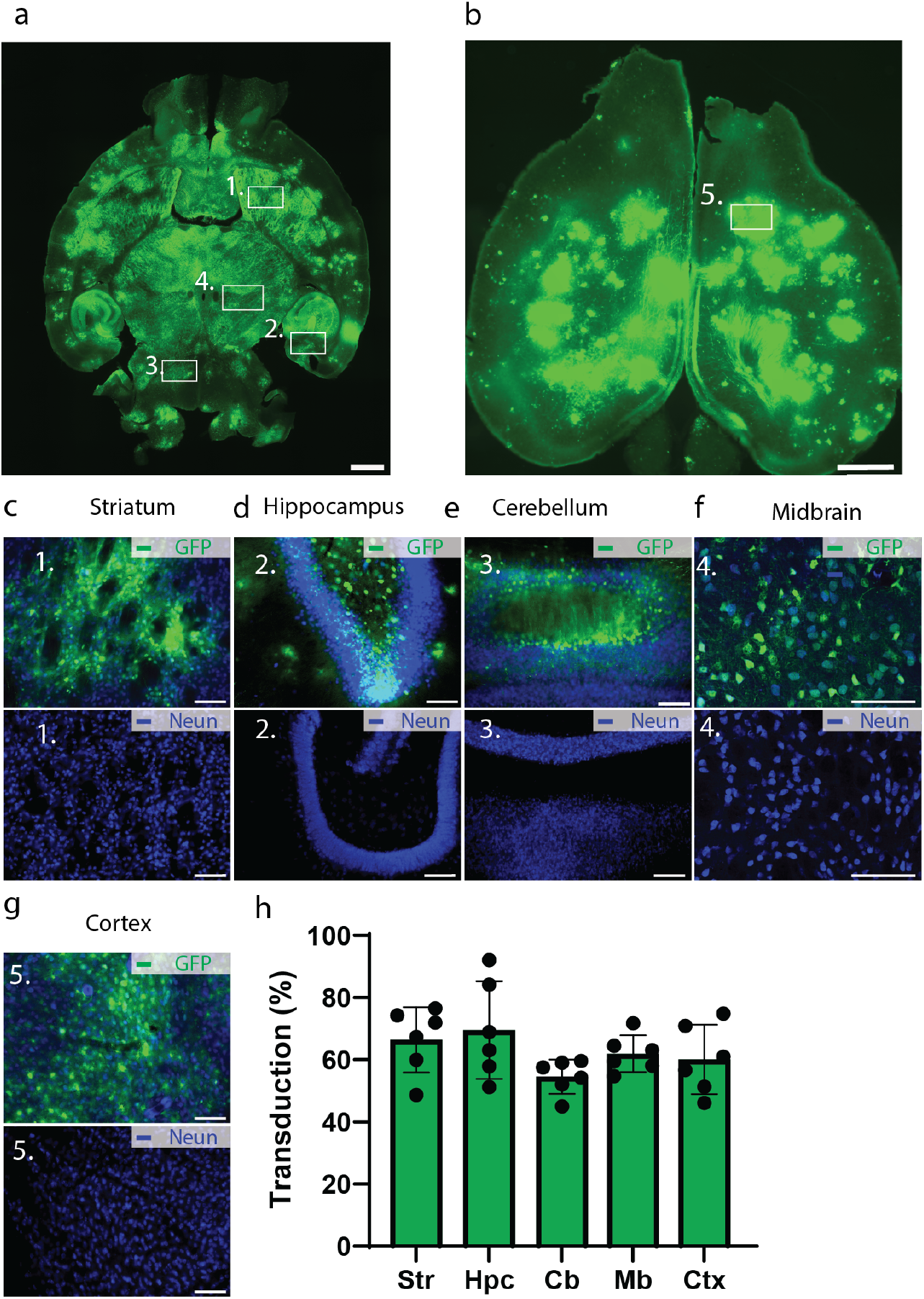
Widespread gene delivery with FUS-BBBO. Immunohistochemistry of anti-GFP, 3 weeks after the FUS-BBBO procedure of brain sections showed an increase in the number of GFP-positive cells in all targeted sites. Representative images of mouse brain sections treated with FUS-BBBO and control mice in Striatum (c), Hippocampus (d), Cerebellum (e), Midbrain (f), and Cortex (g). GFP-transduced cells in green are shown among the neuron cells stained in blue. The number of GFP-positive cells was found to be 66.4±8.4% in the striatum (1), 69.5±12.6 % in the hippocampus (2) 54.5±4.4% in the cerebellum (3) 61.9±4.8% in the midbrain (4), 60.1±8.9% in the cortex (5) (data represents means with 95% CI). There were no significant differences between the transduction efficiency in these regions [p=0.18, F(4,25)=1.713, One-way ANOVA]. Scale bars represent 1000 microns for a), b), and 100 microns, for other panels. (FUS-BBBO, Focused Ultrasound Blood-Brain Barrier Opening; green fluorescent protein, GFP; Analysis of variance, ANOVA)

We then developed a detailed map of transduction efficiency to guide the targeting of FUS-BBBO in various brain regions. To achieve this, we analyzed each targeted site individually and reported transduction efficiency on a site-per-site basis (**Supplementary Fig. 3**). We used a site in the caudate putamen (CPu, site #16, Supplementary Fig. 2c) as a reference for transduction efficiency, since several previous FUS-BBBO studies have targeted Cpu in different species^40–42^. We found that FUS-BBBO was equally effective in the majority of the sites (98 out of 105 sites, or 93.3%, **Supplementary Fig. 3b, d**), with the most prominent difference in the cortex anterior to the striatum and a single site in the hippocampus. The expression across the different targeting depths is provided in **Supplementary Fig. 4**. Overall, these data suggest that FUS-BBBO can lead to effective gene delivery throughout the brain.

### Safety of the FUS-BBBO gene delivery

To evaluate the safety and adverse effects of widespread gene delivery upon FUS-BBBO, we examined the weight loss of mice in all groups. Significant loss of weight would indicate substantial adverse effects of FUS-BBBO on animal well-being, with 15% of the weight loss constituting a humane endpoint. Throughout the treatment, starting from the FUS-BBBO procedure until euthanasia, we weighed the mice twice per week and no significant weight loss was observed in any of the groups (two-way ANOVA with Dunnett’s multiple comparison test, F (3.300, 38.50) = 0.7774, **Fig. 4a**). After 3 weeks of recording the weights, mice in group 2 were euthanized, perfused with neutral buffered formalin (10%), and their brains were extracted for histological analysis. We evaluated neuronal loss and astrocyte activation by analyzing tissue sections for NeuN (a neuronal marker) and glial fibrillary astrocytic protein (GFAP, a marker of astrocyte activation). In this experiment, we focused on group B to allow a direct contralateral control comparison, while allowing the highest possible number of target sites. We counted the neurons in the same group to assess neuronal loss due to FUS-BBBO. We observed no loss of neurons between the targeted sites and contralateral controls (p=0.92 and p=0.78, for group A (n=44 sites across 4 mice) and B (n= 110 sites across 5 mice). respectively, two-tailed heteroscedastic, paired t-test; p=0.63 for group C (n=630 sites across 6 mice in targeted control, and 315 sites across 3 mice for negative control), two-tailed heteroscedastic, unpaired t-test; **Fig. 4c-e**). For the analysis of astrocyte activation, we used positive pixel analysis, as the complex morphology of astrocytes made cell counting more complicated. GFAP-positive pixel analysis showed no significant difference between the targeted and contralateral untargeted sites for groups A and B (p=0.2101, two-tailed heteroscedastic, paired t-test, **Fig. 4f, g).** However, we did observe a difference in GFAP positive pixel counts for the group C, when compared to negative control mice with 6.8±0.4% value for the targeted sites, and 5.9±0.3% for the negative control, constituting 16.9% higher astrocyte activation (data represents means with 95% confidence intervals, p<0.001, two-tailed unpaired heteroscedastic t-test; n=630 sites across 6 mice for the targeted group, and 315 sites across 3 mice for the negative controls, **Fig. 4h**).

**Figure 4.**
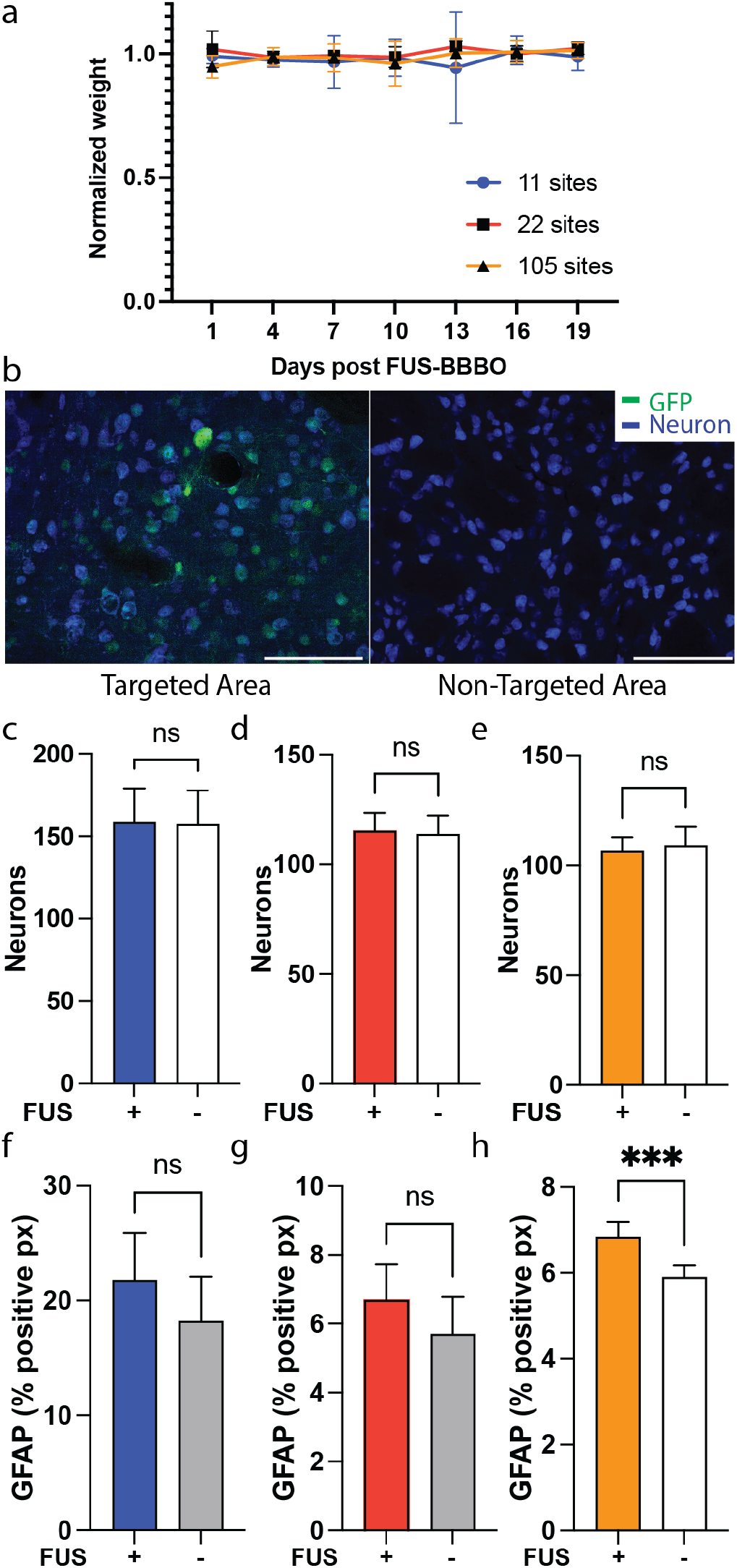
Long-term safety of FUS-BBBO in multiple brain sites. Safety studies 3 weeks post FUS treatment did not show any abnormal weight loss or hemorrhages in mice brains. a) all mice were monitored twice per week for any weight loss or abnormal activity. No significant weight loss was observed. b) Representative images showing neurons (NeuN staining) for the FUS targeted (left) and contralateral untargeted (right) sites. c) Number of neurons was comparable at the sites targeted with FUS and their contralateral controls for group A (p=0.78, n=44 sites per group analyzed in 4 mice; two-tailed paired t-test), and d) in group B (p=0.78, n=110 sites per group analyzed in 5 mice; two-tailed paired t-test). e) Similarly, group C showed no difference in number of neurons in mice targeted with FUS-BBBO or negative control mice that did not undergo FUS-BBBO (p=0.63, 630 sites across 6 mice in the targeted group, or 315 sites across 3 mice in the negative control group, two-tailed unpaired t-test). f) An astrocyte marker, GFAP, was analyzed by positive pixel count in all groups. For group B it showed no significant difference between the number of activated astrocytes (p=0.19, n=44 sites per group analyzed in 4 mice, two-tailed paired t-test), indicating no astrocyte activation at the FUS-targeted sites. g) Similarly, we observed no differences in astrocyte activation in for group B (p=0.21, n=110 per group analyzed in 5 mice, two-tailed heteroscedastic t-test). h) However, we did observe a significant difference in positive pixel counts of GFAP in group C, with 6.8% of positive pixels in FUS-BBBO-targeted group (630 sites analyzed across 6 mice) and 5.9% in the negative control mice (315 sites analyzed across 3 mice, p<0.001, two-tailed unpaired t-test). Scale bars represent 100 microns. (FUS-BBBO, Focused Ultrasound Blood-Brain Barrier Opening; green fluorescent protein.)

### FUS-BBBO widespread CRISPR delivery

Gene editing is a promising technique for treating neurodegenerative disorders. Gene editing agents, such as CRISPR^43^, or Cre-recombinase^44^ can be delivered to specific areas of the brain and have recently been tested through direct intracranial injection^22^ or FUS-BBBO^33,45^. We investigated whether targeting large brain volumes for gene editing would also be feasible with FUS-BBBO. Hence, we used Ai9 transgenic mice^46^, which has a STOP cassette that prevents the expression of the tdTomato gene. We used CRISPR to create a double-stranded break (DSB) in the STOP cassette, which allowed the expression of tdTomato.

We targeted a brain hemisphere with 22 sites in a single insonation session (**Fig. 5a**). We co-delivered two AAV9s encoding SpCas9 under the neuron-specific Mecp2 promoter and the guide RNA (gRNA) under the U6 promoter targeting the STOP cassette. Additionally, the gRNA AAV9 carried GFP under the Synapsin promoter, allowing us to evaluate the efficiency of gene delivery. Activated microbubbles were intravenously injected just before sonication. Three weeks after FUS treatment, mice were sacrificed and brain sections (50um thickness) were stained against tdTomato to examine the efficiency of gene delivery and editing (**Fig. 5b**). We observed 24.1±2.1% transduction efficiency at the targeted sites, compared to significantly lower efficiency of 2.8±1% at the contralateral control sites (p<0.0001, two-tailed paired t-test, numbers represent means with 95% CI, n=110 sites per group in 5 mice, **Fig. 5c**). For comparison, gene editing efficiency, as evaluated by tdTomato expression, was lower – we 7.3±1.4% compared to significantly lower 0.6±0.2% in the contralateral control sites (p<0.0001, two-tailed paired t-test, numbers represent means with 95% CI, n=110 sites per group in 5 mice, **Fig. 5d**). We did not observe significant differences in gene editing in different targeted regions with the tdTomato fluorescence present in 9.8±2.8% of hippocampus and 6.2±3.5% of midbrain and it was 5.2±2.2% of the striatum cells (p=0.13, one-way ANOVA, with Tukey’s post-hoc test, **Supplementary Fig. 5).** To evaluate the safety of such gene editing, we immunostained sections for a neuronal marker (NeuN) to evaluate neuronal loss. We observed a significant loss of neurons with 125.3±9.3 neurons per field of view in the targeted sites, and 152.3±12.9 neurons in contralateral controls (p<0.001, two-tailed paired t-test; n=110 sites in each group for 5 mice).

**Figure 5.**
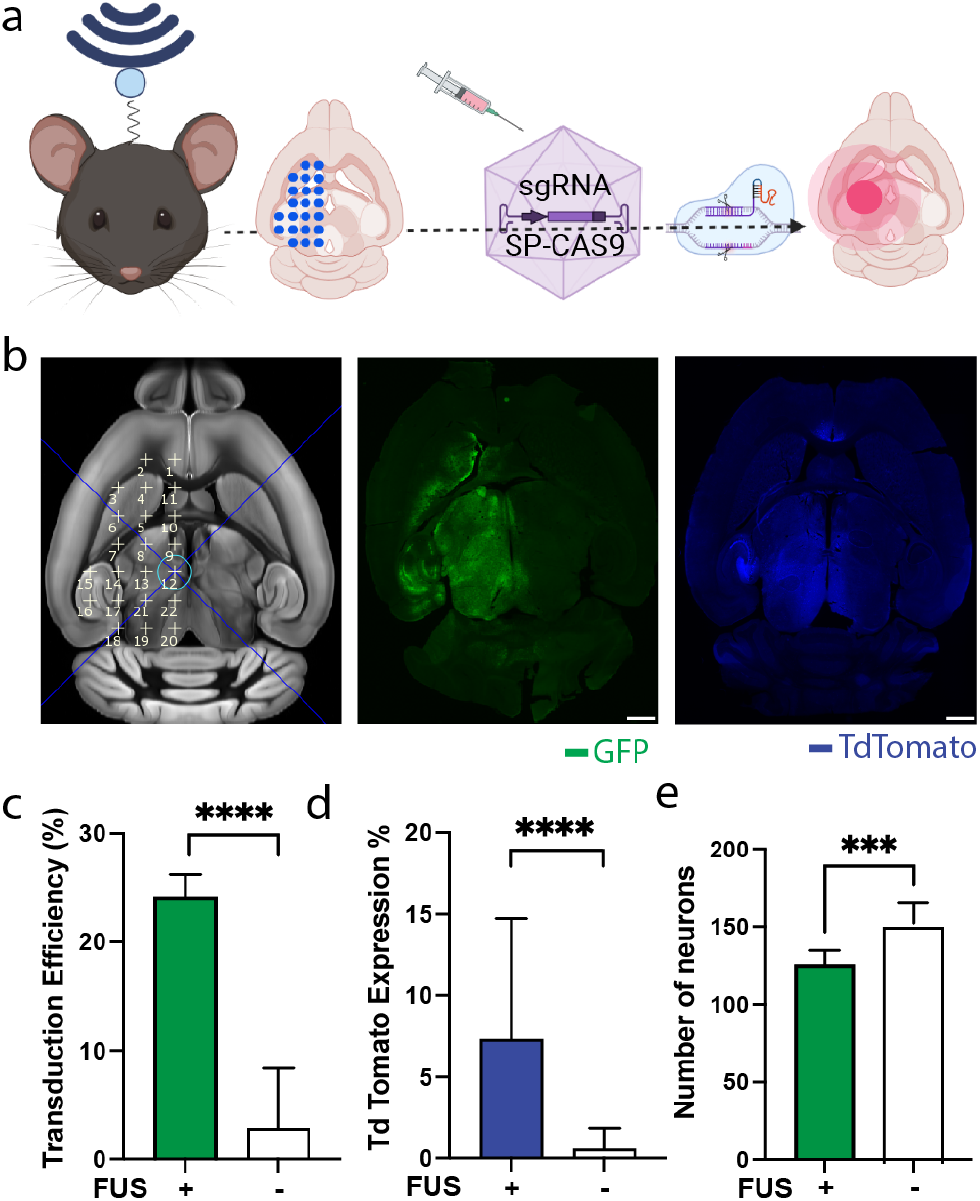
Gene editing after noninvasive gene delivery to multiple brain sites. a) We delivered AAV9 carrying a guide RNA and GFP under neuronal promoter, and another AAV9 carrying spCas9 under MECP2 promoter into 22 sites in the mouse brain. b) We observed GFP and tdTomato expression in the targeted sites, but not in the contralateral controls. c) Transduction efficiency of gRNA and GFP reached 24.1±2.1% and was significantly higher than in untargeted sites (p<0.0001, two-tailed paired t-test, n=110 sites in 5 mice). d) The gene editing efficiency was lower at 7.3±1.4%, but also significantly higher than the untargeted brain regions (p<0.0001, two-tailed paired t-test, n=110 sites in 5 mice). e) We counted neurons at the targeted sites and contralateral controls, and found lower numbers of neurons in the former, indicating a possible neuronal loss. Scale bars represent 1000 microns for a), b).*, p<0.05, **, p<0.01, ***, p<0.001, ****, p<0.0001. (green fluorescent protein, GFP).(Image created with biorender.com)

## DISCUSSION

Although FUS is a technique for targeting specific brain regions with high precision, FUS-BBBO can also be useful in the treatment of disorders affecting large swathes of the brain such as hereditary genetic disorders like Stxbp1 encephalopathy, Huntington’s Disease, or Rett disease^1,5,47^. Additionally, brain regions in nonhuman primates and humans are often larger than the size of the ultrasound beam. For instance, the commonly used research equipment for transcranial FUS-BBBO in humans opens 3 × 6 mm site ^48^, slightly over 0.002% of the brain volume, and 0.63% of e.g. the human hippocampus volume^49^. This suggests that to cover the entire human hippocampus with FUS-BBBO, one would need to perform over 100 FUS insonations.

Moreover, there are many cases in which targeting large swathes of the brain could be particularly beneficial. For example, a large fraction of the hippocampus^50^ or a significant area of the cortex^51^ may need to be targeted using gene therapy to treat temporal lobe epilepsy (TLE). Such regions are infeasible for targeting with direct intraparenchymal injections, which only express genes within a ~2 mm radius from the injection ^52^. Consequently, targeting only the orbitofrontal cortex in an NHP requires 100 injections. Similarly, the newly engineered BBB-permeable viral vectors ^53,54^, lack regional precision and thus cannot be used to target specific brain regions.

Herein, we opened different BBB volumes using FUS. Next, we measured transduction efficiency at the target sites. We found an 18–19.1-fold improvement in gene delivery efficiency at FUS sites compared to the contralateral untargeted sites, but a 252.1-fold improvement compared to mice without FUS-BBBO. This suggests that BBB opening increases gene delivery to off-target sites; however, only one of the tested groups (B) showed significant improvement over control mice that did not receive FUS insonation. Importantly, the transduction efficiency was higher when more sites were targeted. For 11 sites, it was 32.92%; for 22 sites, it was 44.99%, and for 105 sites, it was 60.82%, of average transduction efficiency at the FUS-targeted site (p<0.0001, ANOVA). We found that the CAG promoter was effective for gene delivery, showing expression at each target site. In terms of safety, we found no neuronal loss at the targeted sites after the delivery of GFP. We also observed no weight loss in any of the tested groups that underwent FUS-BBBO, suggesting that FUS-BBBO was well tolerated, even when a large fraction of the brain was targeted. We observed no GFAP accumulation, suggesting that FUS-BBBO did not lead to astrocyte activation. Importantly, it is well validated that FUS-BBBO can cause microglial activation^55–57^; however, that activation is temporary^56^ and can be abrogated by injection of antiinflammatory corticosteroids ^58^.

We also demonstrated that FUS-BBBO delivery can affect gene editing within the brain. Wherein we delivered Spcas9 encoded within AAV9 using FUS-BBBO, targeting 22 sites within the brain. We used the MECP2 promoter, which preferentially targets neurons ^25^ and found that targeted regions had 13.1-fold higher gene editing in neurons in FUS-BBBO targeted sites. We found no significant regional dependence on gene editing efficiency. The overall efficiency of editing over the 21 days of the experiment and at the doses we chose was 7.53%. This efficiency may be sufficient for the treatment of some diseases; for example, when a small number of neurons can produce an enzyme that diffuses throughout the parenchyma^59^. However, disorders targeting the activity of neural circuits ^60^ may require higher transduction efficiency^61^. The efficiency of gene editing via AAV-mediated delivery to the brain is controlled by multiple factors. For example, in current technology, the SpCas9 system is packaged in two AAV vectors, one encoding SpCas9 and the other encoding the guide RNA. Additionally, the large size of SpCas9 necessitates the use of a short promoter, such as MECP2, which is less efficient than some other commonly used promoters, such as CaG^62,63^. Lastly, and possibly more importantly, we found that SpCas9 may be toxic to neurons since we observed that its delivery led to a neuronal loss at the targeted site. This was not observed when we delivered GFP to the same brain sites using the same ultrasound parameters. A similar report suggested toxicity for a different route of delivery of SpCas9 delivery^64^, validating the neurotoxicity of SpCas9, removing the confound of local injury to the tissue causing neuronal loss. The development of new SpCas9 variants that are smaller and may allow the introduction of more efficient promoters or the use of a single AAV capsid encoding both the guide RNA and gene editing enzyme may lead to an improvement in gene editing efficiency and safety. However, to date, we have demonstrated the feasibility of CRISPR gene editing following noninvasive FUS-BBBO delivery to large brain regions.

Overall, we showed the feasibility and safety of opening the majority of the brain volume with FUS-BBBO for gene delivery. We demonstrated that transduction efficiency improves when a larger number of sites are targeted. We also developed a detailed 105-site map of gene delivery efficiency in the mouse brain and showed that this efficiency is remarkable, even across different brain regions. Our protocol uses only 30 s of insonation for AAV delivery, as compared to 120 s in other studies^33,40,65,66^, allowing for faster administration of the protocol, which is important when hundreds of sites are being targeted. Finally, we showed that our optimized FUS-BBBO protocol can be used to affect gene editing within the brain following minimally invasive administration of AAVs. Our data can help benchmark gene efficiency in different brain regions and demonstrate the applicability of FUS-BBBO for potential gene therapy treatment of brain disorders affecting large or multiple brain regions.

## MATERIALS AND METHODS

### Animal subjects

Twenty-four wild-type C57BL/6J mice aged 12 weeks and five Ai9 mice (strain no. 007909) aged 8 weeks were used in this study. Mice were purchased from Jackson Laboratory (Bar Harbor, ME, USA). Animal experiments were conducted in accordance with NIH guidelines and approved by the Institutional Animal Care and Use Committee of Rice University.

### FUS-BBBBO

C57BL/6J male, 12-week-old mice were anesthetized using 2% isoflurane in the air. The fur on their heads was removed using a trimer. Next, a premade catheter washed with heparin and 0.9% saline was placed in the lateral tail vein and secured with tissue glue. Subsequently, the mice were placed on a stereotactic instrument and their heads were fixed with ear bars. The incision area was washed thrice with chlorhexidine scrub, followed by chlorhexidine solution. The mice were then injected via the tail vein with either Evans blue dye or AAV9 carrying CAG-NLS-GFP (1E^10^ viral particles per gram of body weight). The plasmid was a generous gift from Viviana Gradinaru (Caltech, Pasadena, CA, USA) and was obtained from Addgene (plasmid # 104061). Immediately after intravenous injection, mice were injected with approximately 1.5×10^6^ DEFINITY microbubbles (MB) (Lantheus) dissolved in sterile saline (20ul of MB in 980 μL of saline, per gram of body weight)^33^. Within 30 s, the mice were insonated using a 1.53 MHz center-frequency transducer with a full-width half-maximum pressure of ~ 1 × 5 mm, coupled to the head via Aquasonic gel (Parker labs, Fairfield, NJ) placed on the top of the animal’s head. The focal distance was adjusted electronically using software to target specific brain regions. The ultrasound parameters used were as follows: 1.53 MHz, 1% duty cycle and 1 Hz pulse repetition frequency for 30 pulses. Using custom software, we enabled the RK-50 FUS instrument to target multiple regions in a single session. The mice were distributed into three groups A, B and C. The brain was targeted incrementally at 11, 22, and 105 sites for groups A, B, and C, respectively. A single hemisphere was targeted in groups A and B and the cerebellum was excluded, whereas group C had targets throughout the brain. After FUS-BBBO, the incision was closed and the mice were placed in the home cage for recovery. The mice were monitored, and their weights were recorded for three days following surgery and then every three days until day 19.

### Microbubbles

Definity (Perflutren Lipid Microsphere) ultrasound contrast agent was purchased from Lantheus Medical Imaging (Billerica, MA, USA). Based on supplier specifications, a single vial containing up to 1.2 × 10^10^ perflutren lipid microspheres was activated. Therefore, we approximated 1 × 10^10^ microbubbles/1500 μL bubble agent/vial for all calculations.

### CRISPR Delivery

To conduct CRISPR-mediated gene editing in the mouse brains, two transfer plasmids encoding SpCas9 and sgRNA were prepared separately. The plasmid PX551 (Addgene #60957) was used for the SpCas9, expressed under the Mecp2 promoter. To construct the sgRNA plasmid, PX552 (Addgene #60958) was digested with SapI (New England Biolabs, Ipswich, MA) and inserted with a 20-nucleotide spacer sequence (5’-aagtaaaacctctacaaatg-3’), which targets the stop cassette upstream of the tdTomato gene in Ai9 mice. To produce AAV9S, HEK293T cells (ATCC) were transfected with AAV9 Rep-Cap, pHelper, and transfer plasmids and harvested 4 days post-transfection. Cells were lysed using the freeze-thaw method and centrifuged to obtain a clarified lysate containing the viruses. The lysate was loaded into a Quick-seal tube (Beckman Coulter) with the iodixanol gradients of 60%, 40%, 25%, and 15% and subjected to ultracentrifugation to extract the 40% iodixanol layer. The resulting purified AAV9s were washed with PBS and concentrated using the Amicon centrifugal filter unit (MilliporeSigma, Burlington, MA, USA) with a 100 kDa cutoff. The titers of AAVs were calculated by real-time quantitative PCR using SYBR Green PCR Master Mix (Thermo Fisher Scientific, Waltham, MA). FUS-BBBO was performed on Ai9 mice to deliver AAV9s to the left hemisphere with 22 target sites, excluding the cortex and cerebellum. Immediately, the two AAV9s encoding SpCas9 and sgRNA were co-injected intravenously into Ai9 mice at a dose of E^10^ VP/g for each AAV. After 3 weeks of incubation, the mice were sacrificed and their brains were analyzed by immunohistochemistry.

### GFP Immunohistochemistry and Gene expression

For immunostaining, mice were perfused with 10% formaldehyde neutral buffer (ThermoFisher) for 15 min. brains were extracted and fixed in 10% formaldehyde for 16 h. Brains were sectioned (axial plane sections) using a vibratome (Leica VT1200S). Sections (50 μm thick) were used for immunostaining. Sections were first treated with blocking solution for one hour (0.2% Triton X-100, 10% goat serum in 1X phosphate-buffered saline [PBS]) and incubated with the primary antibody (rabbit anti-green fluorescent) (in blocking solution) overnight at 4°C. Sections were washed thrice for 10 min with 1X PBS and incubated in the secondary antibody at room temperature for 4 h. Finally, sections were washed three times in 1X PBS and mounted on glass slides in an Antifade Mounting Medium with DAPI (Vectashield, Thermo Fisher, Waltham, MA). Brain sections were imaged using a Keyence BZ-X810 fluorescence microscope (Osaka, Japan) in the green channel to check for the GFP protein.

### Immunohistochemistry to evaluate the neuronal loss and astrocyte activation

Sections (50 μm thick) were used for immunohistofluorescence. Sections were first treated with blocking solution for one hour (0.2% Triton X-100, 10% goat serum in 1X PBS) and incubated with the primary antibodies anti-Neun (Novus Biologicals, Centennial, CO), anti-GFAP (G3893-Sigma-Aldrich) in blocking solution overnight at 4°C. Sections were washed thrice for 10 min with 1X PBS and incubated with the Alexa 594 and 647 secondary antibodies (Life technologies, Carlsvad, Ca) at room temperature for 4 hr. Finally, the sections were washed three times in 1X PBS and mounted on glass slides in an Antifade Mounting Medium (Vectashield I800, Thermo Fischer, USA). Brain sections were imaged using a Keyence (Osaka, Japan) BZ-X810 fluorescence microscope to check for transduced neurons and astrocytes.

### Histological analysis

To quantify marker proteins, transduced cells were counted at each target site using ZEN 3.3 (Blue edition) software. All images were counted under uniform bright exposure and contrast. The cells in contralateral sites were regarded as a negative control for groups A and B, and cells for mice injected IV with AAV9 carrying GFP, but without FUS-BBBO were negative controls for group C.

For NeuN immune quantification, we used ZEN 3.3 (Blue edition) software. For all image thresholds, a region of interest (ROI) was plotted on the image according to the brain structures in the Allen Mouse Brain Atlas. The number of particles in the ROI was counted separately for each region. The exposure times, image processing, and merging were performed using the same parameters for each experiment. 3 sections per mouse, with an intersectional space of 1000 μm, were stained and quantified for transduction efficiency. The efficiency was averaged for each targeted site for the sections that contained that site.

For GFAP analysis, we used the ImageJ software to analyze the positive pixel count for all targeted sites.

For transduction efficiency analysis DAPI was used to determine the number of neurons in the targeted area, while GFP-positive immunostaining was used to count the neurons. The transduction efficiency was defined as the ratio of the number of GFP-positive cells to the total number of cells. Three sections were analyzed for each mouse. Blinding was not possible since the difference in transduction efficiency was obvious to the naked eye.

To account for the bias and to validate our counting approach, all sites of NeuN staining of all Ai9 mice were counted by two independent blinded scorers (J.C.Y, V.S.). The inter-experimenter variability for the group was 1.7% (126.4±12.2 vs 124.2±12.1 cells on average per image) for the targeted sites, and 5.9% for the contralateral sites (156.6±22.5 vs 147.9±12 cells on average per image).

### Statistical analyses

Comparisons between the two groups were made using a twotailed heteroscedastic t-test. We used a paired t-test if the samples were matched for ipsilateral and contralateral sites and unpaired tests for all other cases. For more than two groups, we used one-way analysis of variance (ANOVA) if a single variable was varied or two-way ANOVA if there were two independent variables. For one-way ANOVA we used the Tukey post-hoc test, and for two-way ANOVA, Dunnett’s multiple comparison test was used. Statistical significance was set at p < 0.05.

## Data availability

The authors declare that all data supporting the results in this study are available within the paper and its Supplementary Information. The raw and analyzed datasets are available from the corresponding author upon reasonable request.

## Acknowledgments

The authors thank Dr. Mingshan Xue (Baylor College of Medicine) for helpful discussions, and Gang Bao’s laboratory (Rice University) for providing the usage of the ultracentrifuge. This research was supported by the Dunn foundation, the related work was funded by The Welch Foundation and Harold Y and G. Leila Mathers foundation.

## Author contributions

JOS and SN designed the experiments, SN and SL performed the in vivo studies, JOS and SN wrote the manuscript, SN, VAS and JY performed histological processing, imaging and counting, HCD and SN performed preliminary in vivo experiments, JOS supervised the study.

## Funding

The work was funded by John S. Dunn Foundation award for collaborative research and by The Welch Foundation grant (C-2048-20200401). Related work was supported by and Harold Y and G. Leila Mathers Foundation.

## Competing interests

The authors declare that they have no other competing interest.

## Notes

### Competing Interest Statement

The authors have declared no competing interest.

