## Supplementary figures for "Acoustically Targeted Noninvasive Gene Therapy in Large Brain Regions"

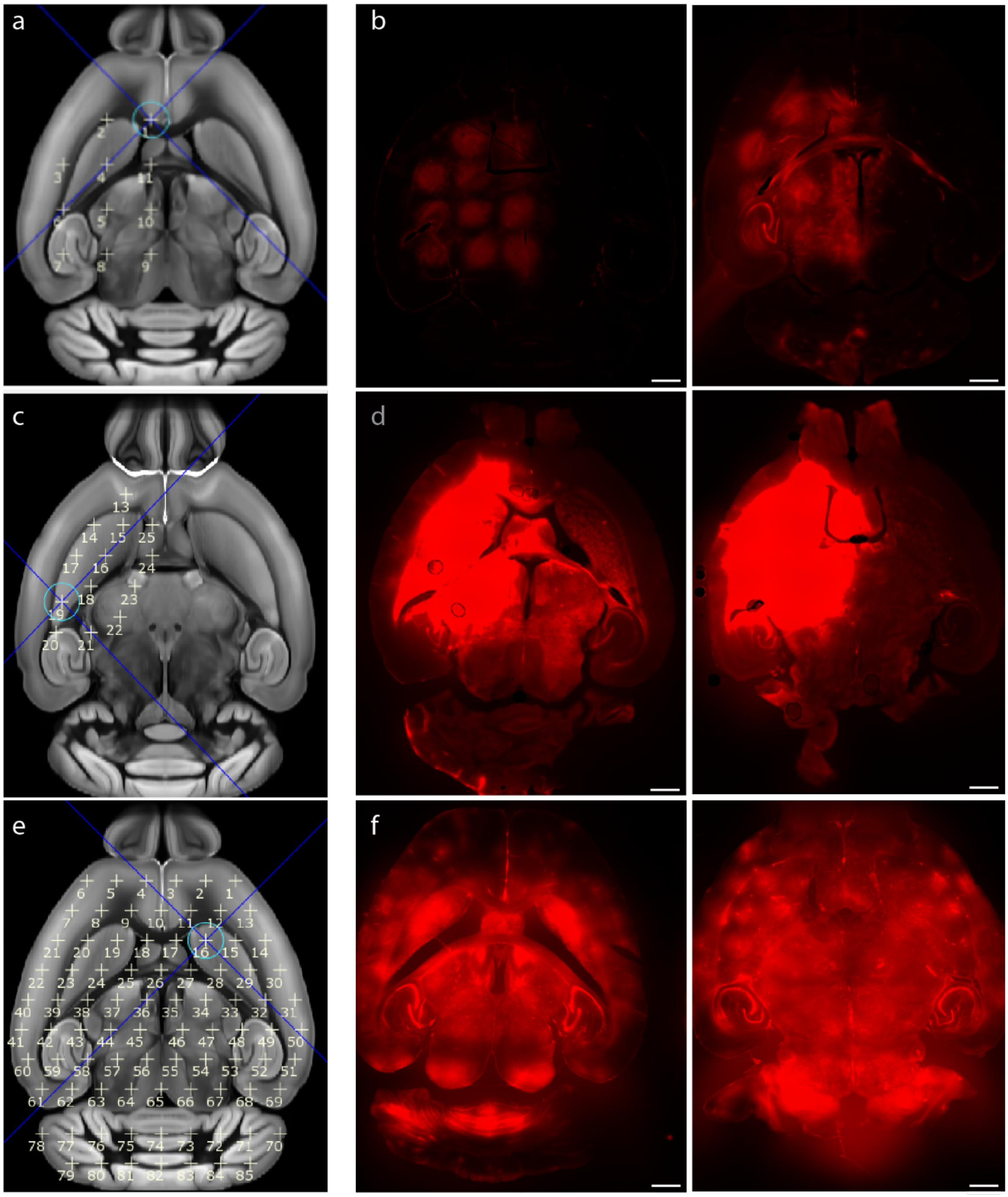


**Supplementary Fig. 1: Delivery of Evans blue dye with FUS-BBBO.** Two mice were tested for each group, with 11, 22, or 85 sites. a) Targeting plan showing the location of the 11 targeted sites. b) EBD fluorescence and bright field image of both tested mice. c) Targeting plan for the 22-site BBB opening, and d) EBD fluorescence for both tested mice. e) Targeting plan showing the location of 85 sites, and f) images of EBD fluorescence for both mice. We have observed no apparent tissue damage in either of the groups. Scale bars are 1000 microns

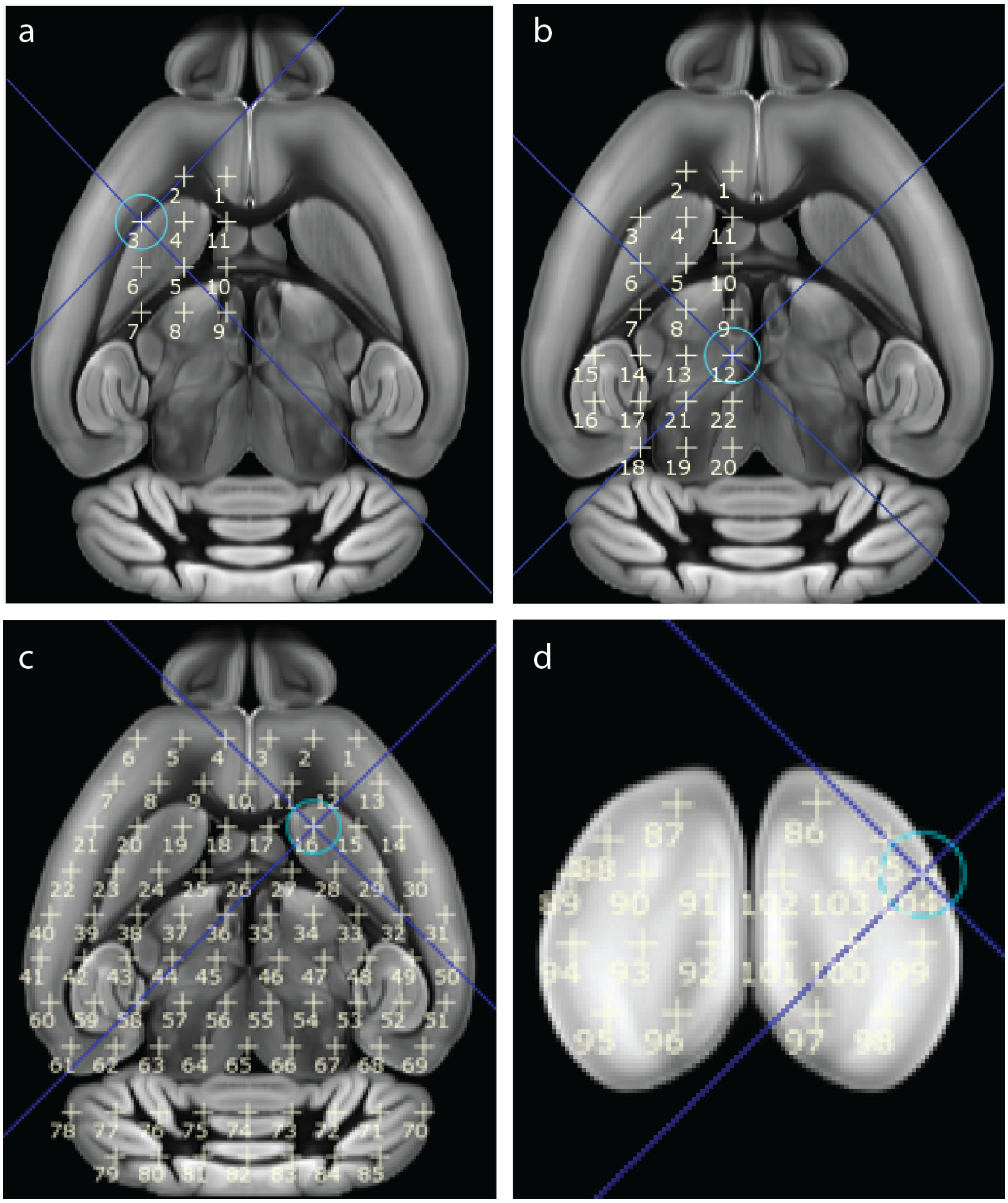


**Supplementary Fig. 2: Targeting strategy for gene delivery**. Site numbers and groups are shown on the axial images of the brain for a) group A, b) group B, and c) group C.

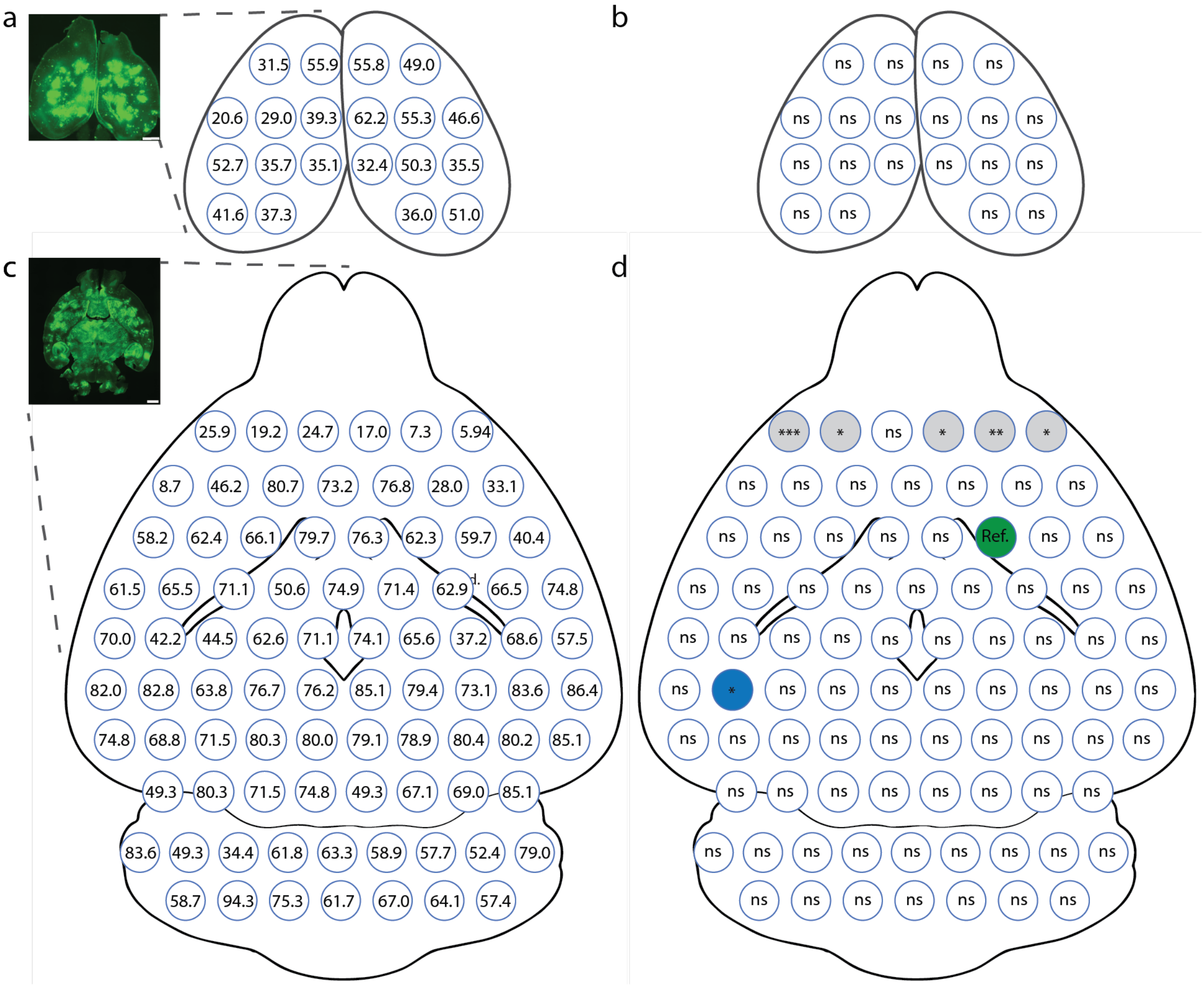


**Supplementary Fig. 3: Transduction efficiency in each of the 105 FUS-BBBO-targeted sites.** a) Representative image of the gene expression in the cortex overlaying hippocampus in group C mice (20 sites), along with the mean transduction efficiency (n=6 mice). Each circle represents a targeted site, and the number inside represents mean transduction efficiency. b) All sites in the cortex are not significantly different than a site transduced in caudate putamen (Cpu), a commonly targeted region in FUS-BBBO studies. Each circle represents a targeted site, and the 'ns' refers to a p-value above 0.05. c) Representative image and the mean transduction efficiency in each targeted site. Each circle represents a targeted site, and the number inside represents mean transduction efficiency. d) Several sites were significantly different than the reference ("Ref.", green) site in Cpu). Most of the sites were at the top targeted row and had lower transduction than Cpu (grey color). However, one site in hippocampus showed higher transduction (blue). ns = p>0.05, * = p<0.05, ** = p<0.01, *** = p<0.001. Scale bars are 1000 microns

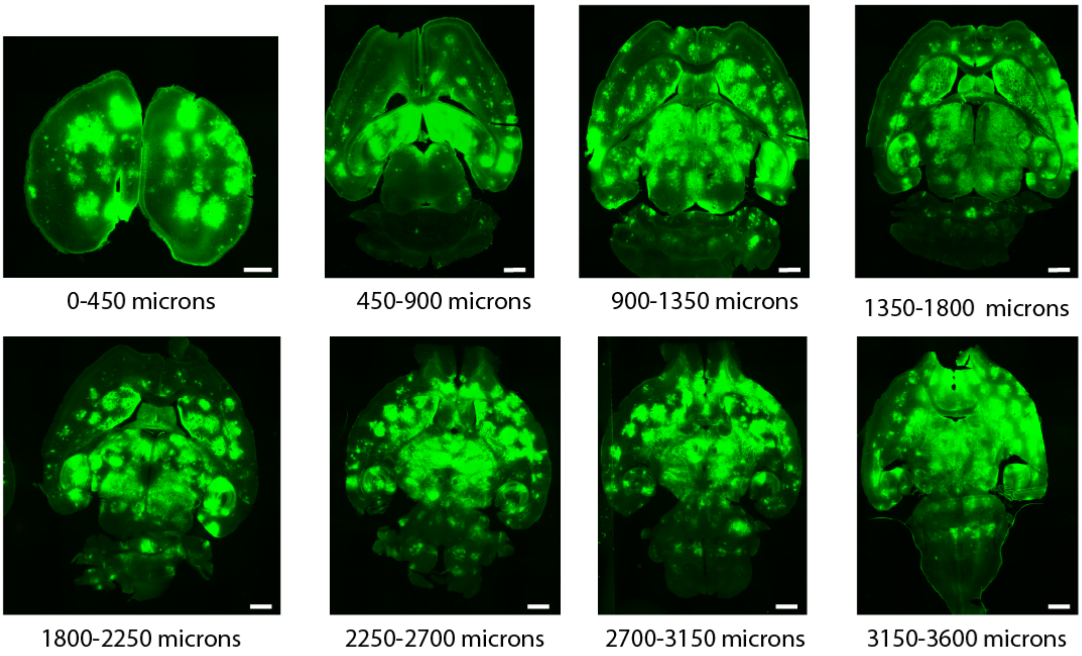


**Supplementary Figure 4. Distribution of gene expression at different depths of targeting.** First image on the left shows the approximate depth of the section from the top of the brain. The 50-micron sections were taken randomly from bins of 450 microns to evaluate how FUS-BBBO enhanced AAV expression changes in 3-dimensions. We observe gene expression at all tested depths, throughout the brain.

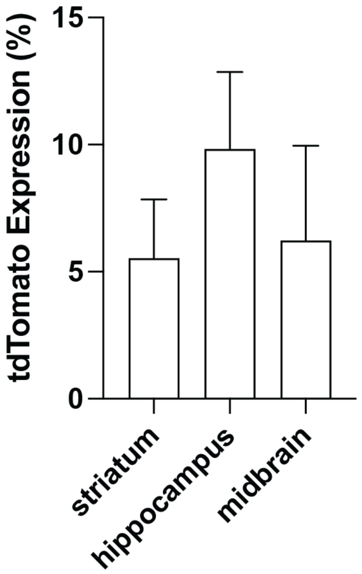


**Supplementary Figure 5.** Gene editing efficiency does not vary between the regions after FUS-BBBO delivery to Ai9 nice. AAV9 carrying SpCas9 under the neuron-specific Mecp2 promoter, and the guide RNA (gRNA) under the U6 promoter targeting the STOP cassette and a GFP under synapsin was injected intravenously and FUS-BBBO applied in the same parameters as in the group B throughout the study (22 sites). We allowed 3 weeks of expression and observed no significant differences in the genome editing efficiency, as evidenced by tdTomato expression among the tested brain regions [one-way ANOVA, with Tukey's post-hoc test, $F(2, 62)= 0.2331$].
